# Uncertainty-aware localization microscopy by variational diffusion

**DOI:** 10.64898/2026.05.01.722206

**Authors:** Clayton Seitz, Jing Liu

## Abstract

Fast extraction of physically relevant information from images using deep neural networks has led to significant advances in fluorescence microscopy and its application to the study of biological systems. For example, the application of deep networks for kernel density (KD) estimation in single-molecule localization microscopy (SMLM) has accelerated super-resolution imaging of densely labeled structures in the cell. However, localization of fluorescent molecules in dense images is a difficult inverse problem with potentially multiple solutions. To model a probability distribution of solutions to this problem, we propose a generative modeling framework for KD estimation in SMLM based on a conditional variational diffusion model (CVDM). In this framework, CVDM is trained to perform localization tasks on low-resolution measurements by modeling a distribution of high-resolution KD estimates. This approach allows us to probe the structure of the distribution on KD estimates and express uncertainty, which is not currently offered by existing deep models for localization microscopy. We demonstrate that this model permits high-fidelity super-resolution, enables the uncertainty estimation of regressed KD estimates, and has important implications for image restoration in single-molecule and super resolution microscopy.

## INTRODUCTION

The fluorescence microscope is an essential tool in the life sciences, enabling interrogation of the spatial structure of biological samples at unprecedented resolution. Recent hardware developments in super-resolution imaging such as structured-illumination microscopy^1^ or stimulated emission and depletion microscopy^2^ have surpassed the classic diffraction limit. However, such super-resolution imaging technologies can require expensive and complex hardware implementations, leading to a significant effort on improving widefield setups and software-based resolution enhancement of microscopy images during postprocessing.

Single-molecule localization microscopy (SMLM) is one super-resolution imaging approach based on widefield microscopy, which has received considerable interest for its relative simplicity. SMLM techniques are a mainstay of modern fluorescence imaging, which localize fluorescent molecules with subpixel accuracy to produce a pointillist representation of the sample at diffraction-unlimited precision^3,4^. Unfortunately, localization can become challenging when fluorescent emitters are dense within the field of view and fluorescent signals significantly overlap^5–8^. Convolutional neural networks (CNNs) have recently been proven to be powerful tools in this context and can be used to extract parameters describing fluorescent emitters such as location, color, emitter orientation, z-coordinate, and background signal^9–11^. For localization tasks, CNNs typically employ up-sampling layers to reconstruct Bernoulli probabilities of emitter occupancy^12^ or kernel density (KD) estimates with higher resolution than experimentally measured images^5,6^. KD estimates (y), inferred from low-resolution experimental measurements (x), are a common representation used in SMLM and can be generated directly from molecular coordinates for supervised learning.

While CNNs have proven powerful, localization in dense scenes remains a difficult inverse problem, and a noisy measurement can underdetermine the number of spots and their locations. Traditional CNN-based methods are deterministic and output point estimates, making computation of uncertainty at test time challenging^13,14^. An alternative approach is to model a distribution on fluorophore numbers and their locations θ, to express uncertainty in these quantities. However, the posterior distribution on fluorophore locations has no known analytical form and can be difficult to compute at test time, since (i) θ is of unknown dimension and (ii) θ can be high dimensional and efficient exploration of the parameter space is often intractable. Moreover, the ability to compute distributions over the entire image in parallel, combined with improved visualization, makes image-based modeling an attractive approach. Therefore, the central goal of this paper is to model a conditional distribution *p*(y|x) on the latent KD estimate y, which is of known dimensionality.

To combine the advantages of generative modeling and deep neural networks, we approach localization in dense scenes using a diffusion model^15-19^. More specifically, we infer high-resolution KD estimates from low-resolution images using a conditional variational diffusion model or CVDM^20^ – a novel generative algorithm inspired by recent variational perspectives on diffusion^21-23^. In the remainder of this paper, we introduce a generative model on localization microscopy images *p*(x) to simulate data for training a model *p*(y|x), based on CVDM. The model is then tested on simulated images and experimental images of fluorescently labeled DNA origami nanorulers and microtubules at various labeling densities.

## METHODS

### SIMULATION OF LOCALIZATION MICROSCOPY IMAGES

The central objective of traditional SMLM is to infer a set of molecular coordinates *θ* = (*θ*_*u*_, *θ*_*v*_) ((*u, v*) represents the 2D laboratory coordinate) from measured low-resolution images x. Here, we approach this localization task by training CVDM on simulated pairs (x,y_0_), where y_0_ is the ground truth KD estimate corresponding to x. Statistical models for likelihood-based simulation of x utilize a likelihood on a particular pixel *p*(x_*k*_|*θ*) which can be determined by a two-step process (Figure 1a). Given the coordinates *θ*, a 2D Gaussian point spread function (PSF) is integrated over pixels (Supplementary Equation 1). The distribution on each pixel *p*(x_*k*_|*θ*) is then taken to be a convolution of Poisson and Gaussian distributions due to shot noise and sensor readout noise, respectively (Supplementary Equation 2). A complete description of the simulation design and parameterization for training CVDM in simulated and experimental scenarios is given in the Supplementary Information.

**Figure 1:**
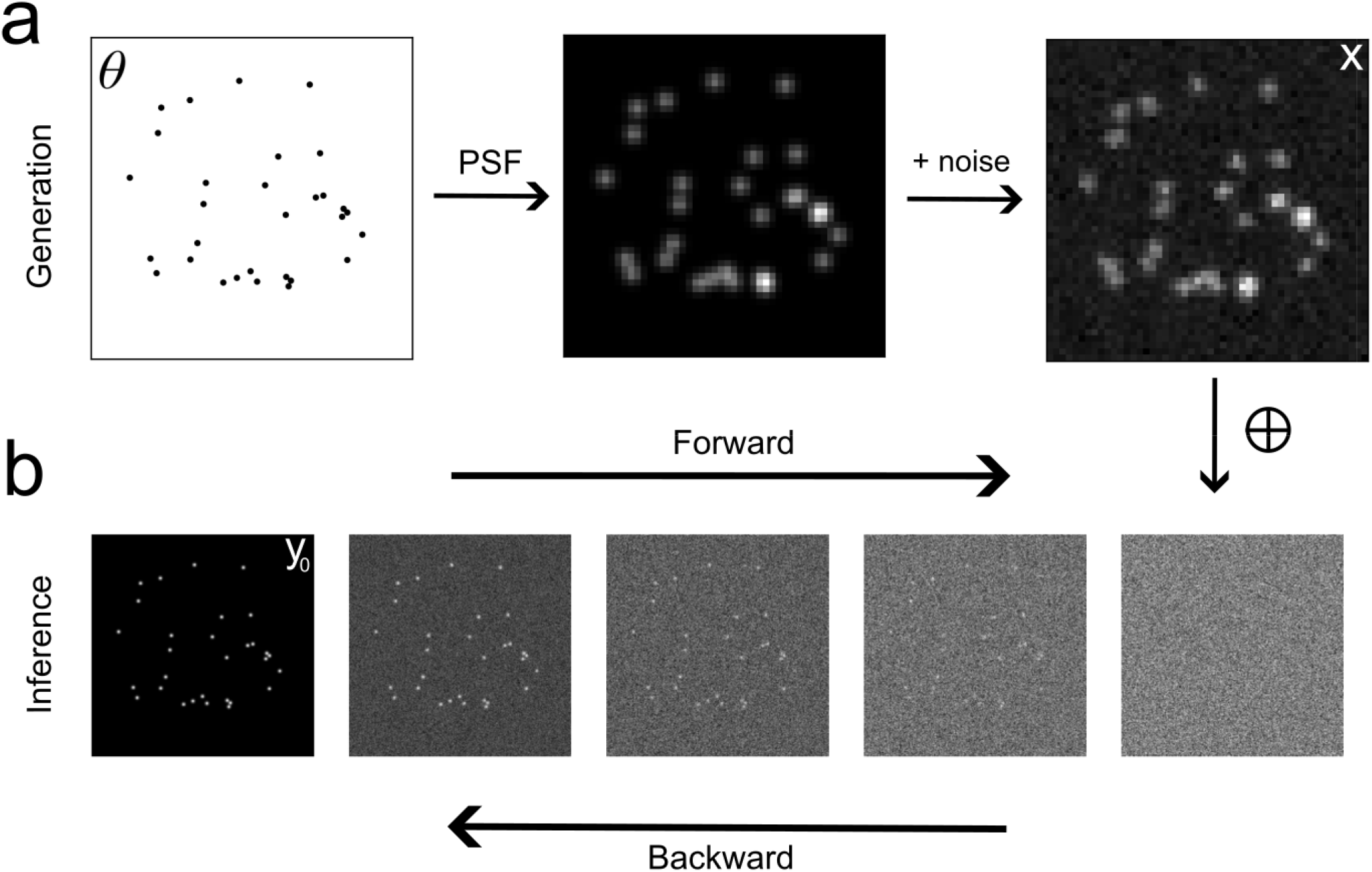
Generation of localization microscopy images and diffusion-based inference to enhance resolution (a) Pipeline of generation of low-resolution localization microscopy image using the point spread function model and noise model. (b) The forward process which transforms the data distribution to pure Gaussian noise and reverse process which involves concatenation of 4x upsampled low-resolution images with pure Gaussian noise and inference of a high-resolution image via the reverse process.

### LOCALIZATION MICROSCOPY BY VARIATIONAL DIFFUSION

In the diffusion framework, we learn a conditional generative model on KD estimates from pairs (x, y_0_) in the simulated training set. For a given y_0_, Gaussian noise is added to y_0_, and we train a deep learning model to iteratively denoise the image, conditioned on the low-resolution x (Figure 1b). Formally, we iteratively destroy structure in y_0_ via a fixed Markov chain referred to as the forward process. Suppose time runs over the continuous interval [0,1] and the diffusion process is sampled at *T* time steps in intervals Δ*t*. We have the following transition distribution in the forward process:

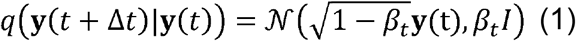

where the value of *β*_t_ controls the variance of the transition of y(*t*) over a period Δ*t*. For inference, the reverse process reads:

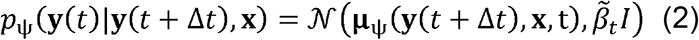

where µ_ψ_ is the expected value of the transition and 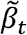 is the variance. Both are computed from y(*t*+ Δ*t*) by the model *ψ*. Additional details of the diffusion model and the training objective are discussed in the Supplementary Information.

### TESTING THE MODEL ON SIMULATED IMAGES

To verify the localizations made by the diffusion model, we simulated low-resolution localization microscopy images with various densities and used CVDM to predict the corresponding KD estimate. The simulated dataset consists of 64×64 low-resolution images x with zero background signal and corresponding 256×256 ground-truth KD estimates. The PSF width in the low-resolution images is σ_x_ = 100nm to match common imaging conditions in super-resolution microscopy. For a KD estimate predicted by the diffusion model, objects are detected using the Laplacian of Gaussian (LoG) detection algorithm and 2D Gaussian fitting, which permits more direct computation of localization error, compared to image similarity measures. Coordinates are obtained by identifying local maxima in the KD estimate. To compute the localization error, a particular LoG localization in the KD estimate is paired to the nearest ground truth localization and is unpaired if a localization is not within 5 KD estimate pixels (125nm) of any ground truth localization. For the ensemble of successfully paired localizations, the localization error is computed as the standard deviation of the Euclidean distance between the ground truth and estimated locations.

### TESTING THE MODEL ON EXPERIMENTAL IMAGES

For testing the CVDM model on experimental data, DNA origami 94 nm nanorulers (GATTA quant) tagged with Alexa555 were immobilized in a coverglass bottom chamber by incubation with 1 mg/mL BSA-Biotin in PBS at room temperature for 5 min followed by washing with PBS. Then, the sample was incubated with 1 mg/mL neutravidin in PBS at room temperature for 5 min. The chamber was then washed with immobilization buffer (PBS supplemented with 10 mM Magnesium Chloride). The sample was then incubated with DNA origamis diluted in immobilization buffer. After incubation, the sample was washed with immobilization buffer. Images of the nanorulers in 2D were collected using a custom Olympus IX83 microscope body equipped with an Olympus 100x 1.49NA oil-immersion objective. Images were projected onto an ORCA-Fusion sCMOS camera (Hamamatsu). The microscope was controlled using Micromanager software. Nanorulers were excited using oblique illumination with a 561 nm laser (Excelitas) held at ~10 mW, as measured at the back focal plane of the objective. When necessary, the gain of the camera used in experiments was measured by fitting a photon transfer curve (PTC) to a movie of uniform background signal, using a known readout noise variance measured by capping the camera sensor. Microtubule data is obtained from a public repository and therefore no experiments were carried out in the present study.

## RESULTS

### LOCALIZATION ERROR FOR SIMULATED IMAGES

Qualitative inspection of CVDM predictions is carried out by comparison of predicted KD estimates with the ground truth (Figure 2a). Quantitatively, we find a density dependence to localization error in the *u* and *v* directions, over the tested signal range (Figure 2b,c). The pairing of detected and ground-truth localizations is also used to quantify the quality of localization recovery. To do this, we measure a density-dependent precision P = TP/(TP+FP) and recall R= TP/(TP+FN) where TP denotes true positive localizations, FP denotes false positive localizations (predicted locations that do not correspond to any ground truth localizations), and FN denotes false negative localizations (ground truth localizations are missing in predicted locations). In ideal conditions, there are no false positives and no false negatives and therefore the precision and recall are both unity. In simulations, we find that precision and recall decay with increasing density, while maintaining strong performance over the density regimes tested (Figure 2d,e).

**Figure 2:**
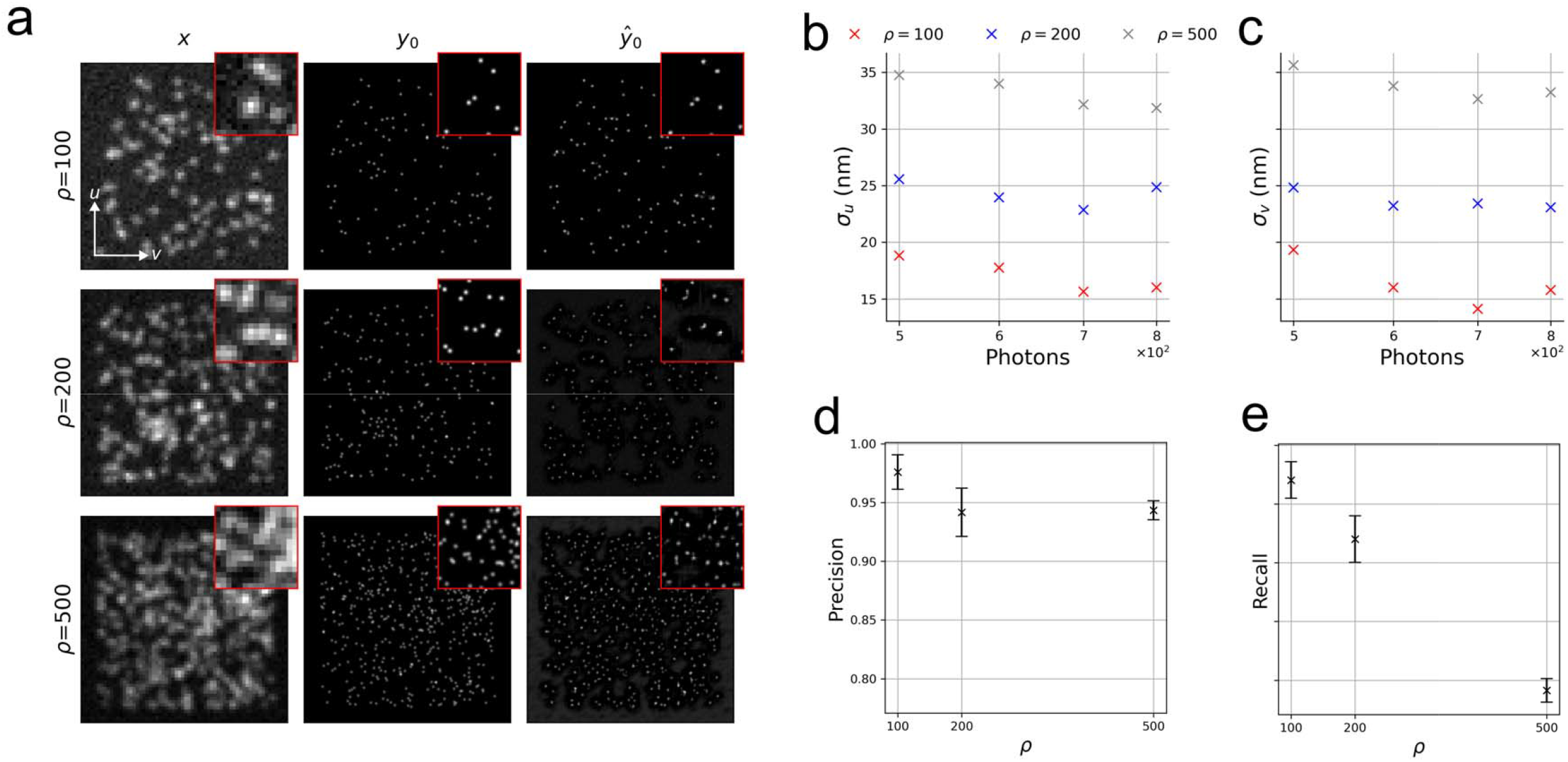
Performance of conditional variational diffusion on simulated data (a) Example low-resolution (left), high-resolution (middle), and estimated high-resolution (right) localization microscopy images at fluorophore numbers in the image (b,c) Localization uncertainty in the u and v directions as a function of incident photon count, for various densities (d,e) Precision and recall of localized spots as a function of. Error bars represent standard deviations over ten batches of data.

### SAMPLE VARIATION FOR SIMULATED IMAGES

Leveraging the ability of a diffusion process to model a distribution on high-resolution KD estimates, as opposed to a deterministic point estimate of y_0_, we assess the sample variation on simulated localization microscopy data. Upon inspection of samples from the diffusion model, we find that computing a pixel-wise average and standard deviation can summarize model predictions (Figure 3a). However, careful inspection of the individual samples can reveal the appearance/disappearance of predicted spots in the KD estimate across samples, expressing uncertainty in the number of spots in the KD estimate (Figure 3b). In other words, the CVDM directly models the joint distribution of the number of fluorescent spots and their respective locations, where each sample is a proposed solution to the image restoration problem. Furthermore, variation in the predicted locations of spots in the KD estimate is apparent, with more variation in higher density samples (Figure 3c).

**Figure 3:**
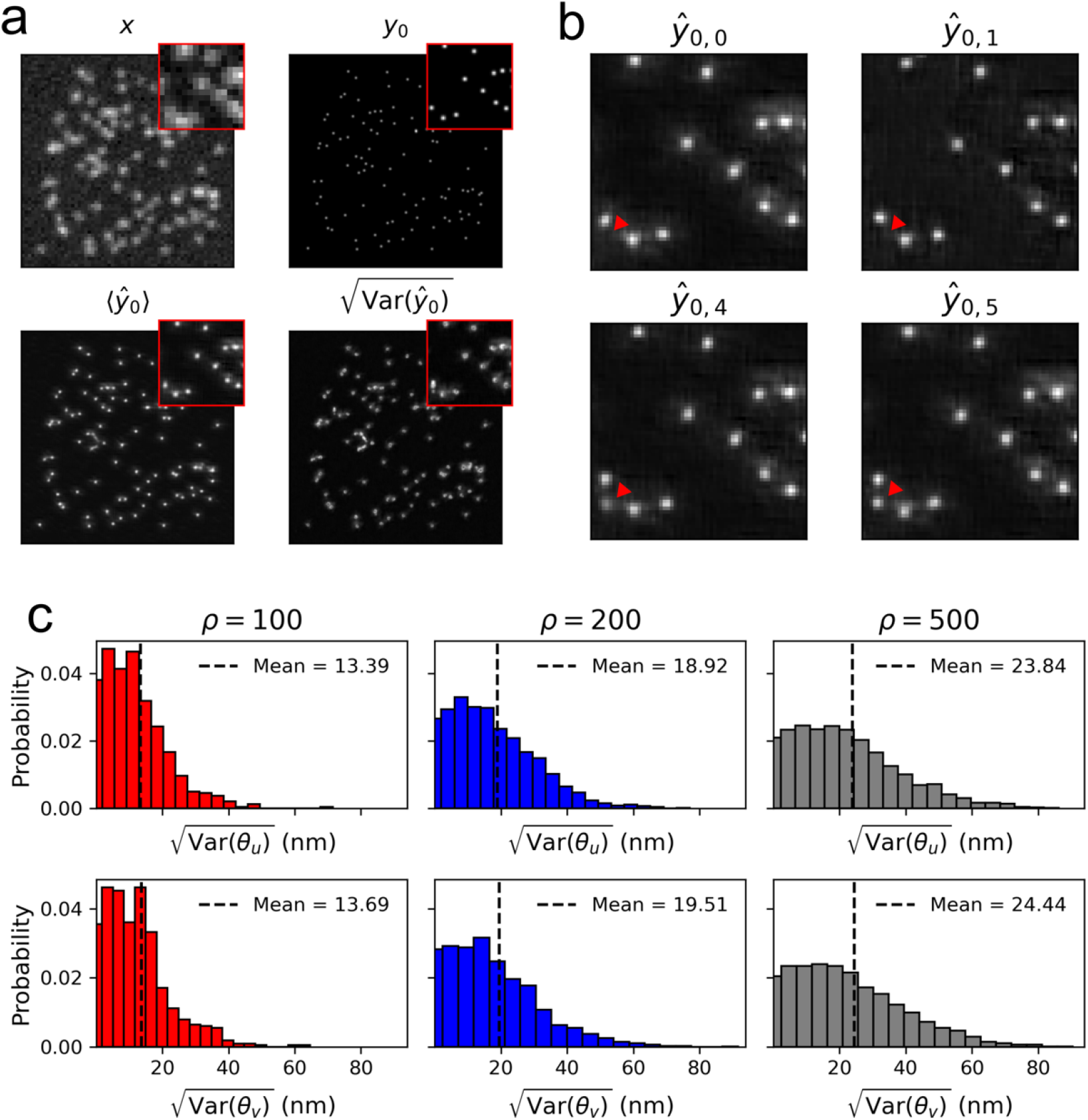
Sample variation for simulated localization microscopy data (a) Low-resolution, high-resolution target, pixel-wise averaged prediction, and pixel-wise standard deviation (b) Samples from the diffusion model on high-resolution images for the inset region shown in (a) (c) Distribution of the standard deviation in emitter locations for various emitter densities. For each density, 3 low-resolution images are simulated and 20 high-resolution KD estimates are sampled using the diffusion model and localizations are matched to ground-truth points to compute the standard deviation over spots.

### SUPER-RESOLUTION OF DNA ORIGAMI NANORULERS

To validate the CVDM model on experimental super-resolution, we train a robust model over a wide variety of simulated images while fixing the experimental PSF and noise characteristics of the camera used. DNA origami nanorulers used for imaging were tagged with Alexa555, with 94nm spacing between marks (Figure 4a). We find that CVDM can predict KD estimates on experimental nanoruler images and pairs of localizations are observed at separations consistent with the known nanoruler spacing (Figure 4b). In addition, we analyze posterior variability for an anisotropic spot by identifying the most probable number of emitters in a region of interest over 100 samples from CVDM, followed by fitting a gaussian mixture model to the localizations with a fixed number of components (Figure 4c).

**Figure 4:**
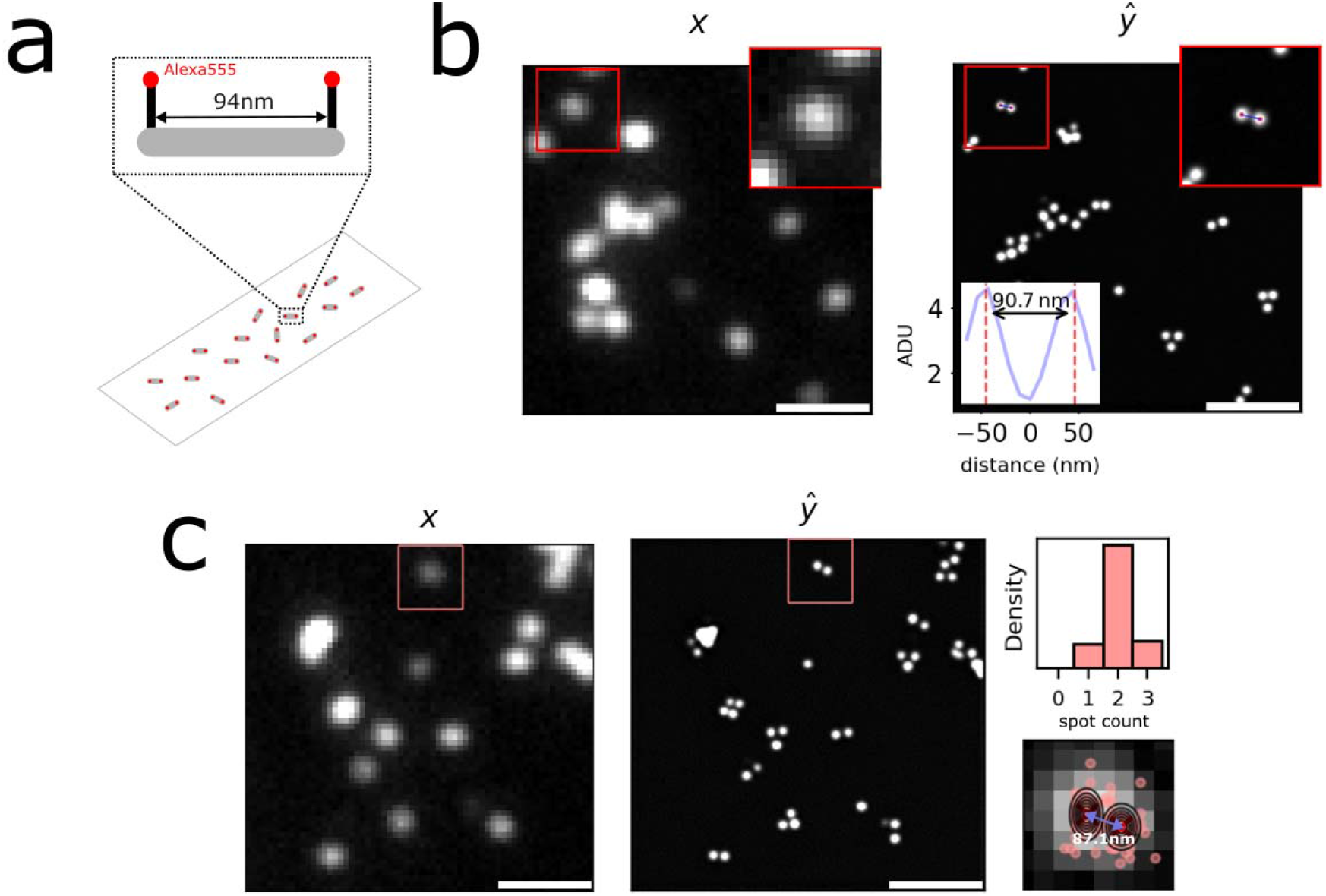
Super-resolution of DNA origami nanorulers using CVDM. (a) Cartoon of a nanoruler coated microscopy slide (b) Low-resolution image of nanorulers and single sample from CVDM with a putative nanoruler (inset). (c) Low-resolution image of nanorulers with multiple samples from CVDM. Anisotropic intensity spot marked in pink. Histogram of the spot count in the pink box and scattered localizations are shown for 100 high-resolution samples from CVDM. Contours show a two-component Gaussian fit with a distance of 87.1nm between components. Scalebars are 500nm.

### SUPER-RESOLUTION OF MICROTUBULES

To further validate the model, we used a single sample from CVDM to improve the resolution of images in a SMLM sequence of microtubules^24^. The dataset consists of images with high labeling density (HD) and moderately dense images constructed by summing five consecutive frames of sparsely labeled microtubules imaged over a long sequence (LS) of 15k frames (Figure 5a). The LS and LS-SUM datasets together can be used to validate the quality of CVDM predictions on moderately dense experimental data, while the HD dataset serves as an additional comparison between multi-emitter localization using the ThunderSTORM^25^ algorithm and CVDM on dense data. By enhancing the resolution of dense scenes with CVDM, we could resolve neighboring fluorophores which were unresolvable in a widefield image (Figure 5b). Furthermore, we compared super-resolution reconstructions obtained by CVDM/ThunderSTORM applied to LS-SUM and ThunderSTORM applied to LS data, as well as ThunderSTORM and CVDM on HD data (Figure 5c). We find strong agreement between all three super-resolution images for LS/LS-SUM (Figure 5d). However, we find that CVDM produces a more detailed super-resolution image for HD data, potentially due to more accurate estimates by CVDM in the dense case (Figure 5e).

**Figure 5:**
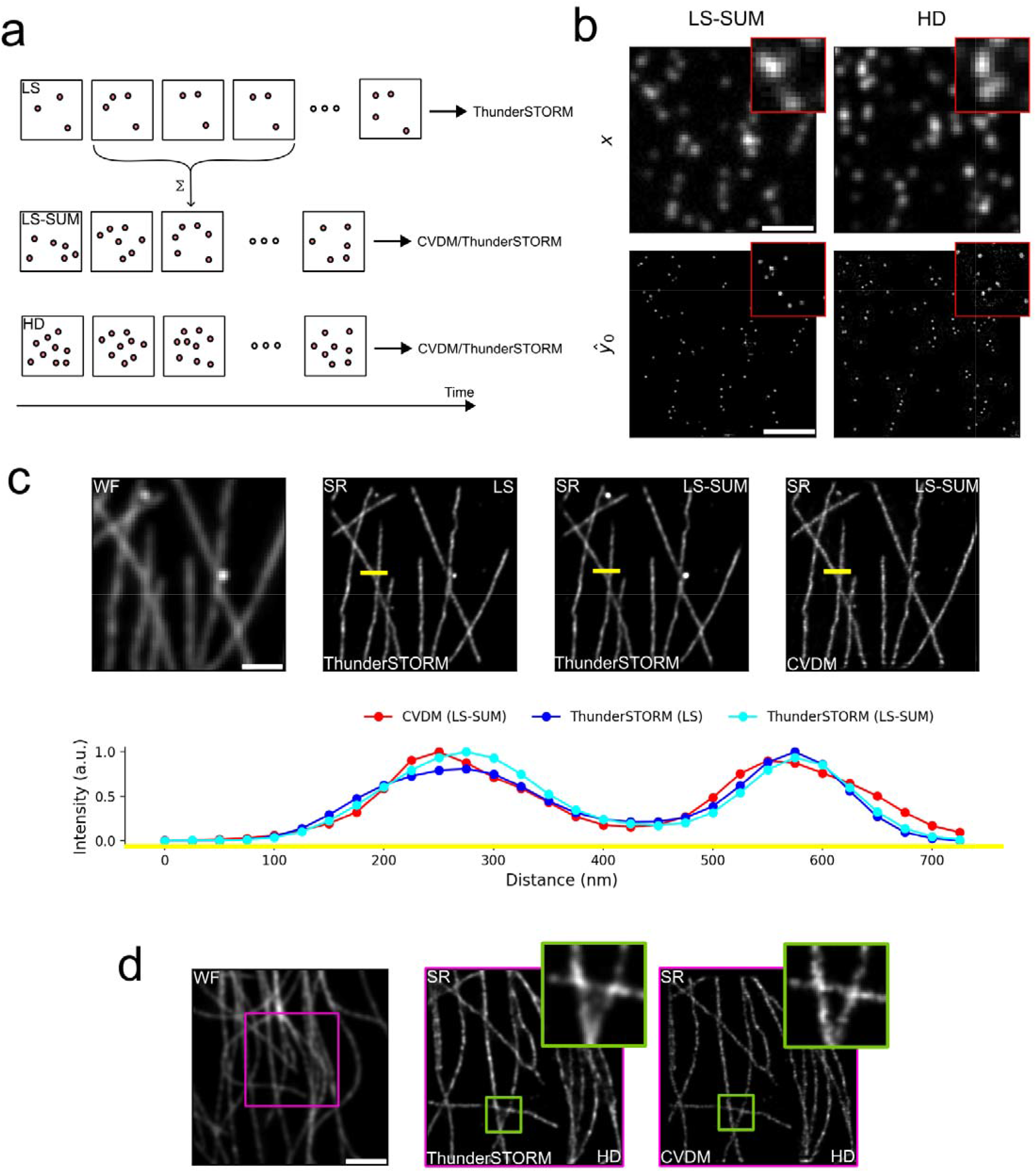
Super resolution of microtubules with CVDM (a) Depiction of long-sequence (LS) of 15k frames, summed long-sequence (LS-SUM) generated by summing five consecutive frames to generate 3k frames and high-density (HD) sequence of 500 frames (b) Example frames from a SMLM sequence and a single corresponding sample from CVDM for LS-SUM and HD data (c) Widefield image of microtubules and super-resolution images found by applying ThunderSTORM and CVDM algorithms to each frame of the LS/LS-SUM sequence. Comparison of the cross-sectional profiles of the super-resolution images produced by CVDM and ThunderSTORM (d) Widefield image of microtubules and super-resolution images generated by applying CVDM and ThunderSTORM to the HD sequence. Inset images are a comparison of CVDM and ThunderSTORM super-resolution images for the selected region. All scalebars approximately 3um.

## DISCUSSION

A central advantage of CVDM is that it reframes dense localization microscopy as a probabilistic image restoration problem rather than a deterministic regression task. In this formulation, multi-emitter localization is performed jointly across the full field of view, avoiding explicit partitioning of the image into local patches or iterative model selection over emitter number. This image-based perspective makes the inferred distribution over spot number and location more accessible for visualization and interpretation, allowing uncertainty to be examined directly in the predicted KD estimates. Rather than producing a single deterministic reconstruction, CVDM generates a distribution of plausible solutions, enabling direct inspection of ambiguity in emitter count and position. In practice, this provides a more natural framework for assigning confidence to localization outcomes in dense scenes, where overlapping signals can admit multiple plausible explanations.

Furthermore, experimental localization microscopy often lacks suitable ground truth data for training. We have approached this issue by simulating a diverse set of training images or approximating experimental labeling density, photon count distribution of individual fluorophores, sensor noise characteristics, and background signal. Experimental validation of the model can also be challenging but tractable if structural information is known apriori. Alternatively, summation of consecutive images to generate high-density data is a suitable approach for low-noise camera sensors.

The principles underlying this method resonate across various fields, suggesting its potential applicability in microscopy beyond localization. Extensions to image denoising and other inverse problems suggest that this framework may provide a general approach for uncertainty-aware modeling across bioimaging and medical imaging applications. Importantly, the utilization of diffusion models for uncertainty estimation aligns with a broader trend in leveraging probabilistic frameworks for enhancing deep learning applications. Diffusion modeling is a growing field, which will benefit from new architectures and techniques for faster sampling. By bridging these interdisciplinary boundaries, this method not only addresses a critical need in localization microscopy but also contributes to the advancement of uncertainty-aware deep learning methodologies.

## CONCLUSIONS

In this work, we applied a conditional variational diffusion model (CVDM) for dense single-molecule localization microscopy and demonstrated its ability to recover high-resolution kernel density estimates from low-resolution measurements. By framing localization as an image-based generative modeling problem, the method enables simultaneous multi-emitter localization across the full field of view while capturing uncertainty in both fluorophore number and position. We showed through simulations that CVDM maintains strong localization precision and recall across a range of emitter densities and validated its performance on experimental nanoruler and microtubule datasets. Importantly, the probabilistic nature of the model allows direct sampling of plausible solutions, providing a principled framework for uncertainty quantification that is difficult to achieve with deterministic approaches. These results suggest that diffusion-based generative models offer a powerful and flexible alternative for super-resolution microscopy, particularly in challenging high-density imaging regimes.

## DECLARATIONS

### Ethics approval and consent to participate

Not applicable

### Consent to publication

Not applicable

### Availability of data and materials

The datasets generated and/or analyzed during the current study are available from the corresponding author on reasonable request. Relevant software can be found at https://github.com/cwseitz/cvdm-smlm

### Author’s contributions

Conceptualization, C.S. and J.L.; methodology, C.S.; software, C.S.; validation, C.S.; formal analysis, C.S. and J.L.; investigation, J.L.; resources, J.L.; writing—original draft preparation, C.S.; writing—review and editing, C.S., and J.L.; visualization, C.S. and J.L.; supervision, J.L.; project administration, J.L.; funding acquisition, J.L.

## Acknowledgements

Not applicable

