## Supplemental Data for "Uncertainty-aware localization microscopy by variational diffusion"

**SIMULATION OF LOCALIZATION MICROSCOPY IMAGES**

Consider an idealized scenario where an isolated fluorescent molecule exists in the image plane. This molecule is imaged by a microscope with point spread function $O\left( u,v \right),$which is presumed to be an isotropic Gaussian in two-dimensions. For a particular pixel $k$of width $\delta$centered at $(u_{k},v_{k})$along the first image dimension, we have

$\Delta E_{u}=\int_{u_{k}-\delta/2}^{u_{k}+\delta/2} O(u)du = \frac{1}{2}\left( \mathrm{erf}\left( \frac{u_{k}+\delta/2-\theta_{u}}{\sqrt{2}\sigma_{\mathbf{x}}} \right)-\mathrm{erf}\left( \frac{u_{k}-\delta/2-\theta_{u}}{\sqrt{2}\sigma_{\mathbf{x}}} \right) \right)$ (1)

The expected number of photons at each pixel is then $\omega_{k}\propto N_{0}\Delta E_{u}\Delta E_{v}$ where $N_{0}$ is a photon count rate per fluorophore. The likelihood of digital units measured at a particular pixel $p(\mathbf{x}_{k}|\theta)$, where $k$ is used as a pixel index, is taken to be a convolution of Poisson and Gaussian distributions, due to shot noise $p(s_{k})=\mathrm{Poisson}(\omega_{k})$ and sensor readout noise $p(\zeta_{k}\mathcal{)=N(}o_{k},w_{k}^{2})$ where $\omega_{k}$ is the signal rate and $o_{k}$ and $w_{k}^{2}$ are the sensor offset and readout noise variance, respectively.

$p\left( \mathbf{x}_{k} | \theta\right)=A\sum_{q=0}^{\infty} \frac{1}{q!}e^{-\omega_{k}}\omega_{k}^{q}\frac{1}{\sqrt{2\pi}w_{k}}e^{-\frac{(\mathbf{x}_{k}-g_{k}q-o_{k})^{2}}{2w_{k}^{2}}}$ (2)

where $A$ is a normalization constant. For the sake of generality, we include a per-pixel gain factor $g_{k}$, which can vary depending on the camera sensor used. Using this, sampling from the convolution distribution in (1) is carried out by $\mathbf{x}_{k}=s_{k}+\zeta_{k}$ for $s_{k}\sim\mathrm{Poisson}(\omega_{k})$ and $\zeta_{k}\mathcal{\sim N(}o_{k},w_{k}^{2})$.

**DIFFUSION MODEL**
The CVDM^20^ model consists of three parts $\psi=(\tau,\phi,\lambda)$ where $\tau$ is a convolutional network which produces a temporal map, while $\lambda$ and $\phi$ are U-Net architectures with skip connections. Model $\lambda$ computes the spatial component of the variance schedule while model $\phi$ is a denoising model used for noise estimation.

During the forward process, the image signal level $\gamma\left( t,\mathbf{x} \right)$ decays exponentially according to $\dot{\gamma}(t,\mathbf{x})=-\beta(t,\mathbf{x})\gamma(t,\mathbf{x})$, thus $\gamma\left( t,\mathbf{x} \right)= e^{-\int_{0}^{t} \beta\left( s \right)\mathrm{ds}}$ where $\beta\left( t,\mathbf{x} \right)$ is an instantaneous noise variance. This noise variance is a product of temporal and spatial parts: $\beta\left( t,\mathbf{x} \right)=\tau(t)\lambda(\mathbf{x}\boldsymbol{)}$. However, in practice, the forward process must occur by integrating over discrete time intervals. The additional variance injected in a time interval $\Delta t$ is defined as $\beta_{t}$

$$q\left( \mathbf{y}\left( t+\Delta t \right)|\mathbf{y}\left( t \right) \right)\mathcal{=N}\left( \sqrt{1-\beta_{t}}\mathbf{y}\left( t \right),\beta_{t}I \right)$$

where $\beta_{t}=1- e^{-\int_{t}^{t+\Delta t} \beta\left( s \right)\mathrm{ds}}= 1-\frac{\gamma\left( t+\Delta t,\mathbf{x} \right)}{\gamma\left( t,\mathbf{x} \right)}$. The stepped reverse process reads:

$p\left( \mathbf{y}\left( t \right)|\mathbf{y}\left( t+\Delta t \right) \right)\mathcal{=N}\left( \boldsymbol{\mu}_{\psi}\left( \mathbf{y}\left( t+\Delta t \right) \right),\tilde{\beta}_{t}I \right);$ $\tilde{\beta}_{t}=$ $\beta_{t}\frac{1-\gamma\left( t,\mathbf{x} \right)}{1-\gamma\left( t+\Delta t,\mathbf{x} \right)}$

where $\boldsymbol{\mu}_{\psi}$is the unknown expected value of the transition$.$To obtain $\boldsymbol{\mu}_{\psi}\left( \mathbf{y}\left( t+\Delta t \right) \right)$notice that by the definition of the forward process $\mathbf{y}\left( t+\Delta t \right)\sim q\left( \mathbf{y}(t+\Delta t)|\mathbf{y}_{0} \right)$ it can be reparametrized as

$$\mathbf{y}\left( t+\Delta t \right)=\sqrt{\gamma\left( t+\Delta t,\mathbf{x} \right)}\mathbf{y}_{0}+\sqrt{1-\gamma\left( t+\Delta t,\mathbf{x} \right)} \boldsymbol{\epsilon}$$

where $\boldsymbol{\epsilon}\sim\mathcal{N}\left( 0,I \right)$. We can estimate $\mathbf{y}_{\mathbf{0}}$ by rearranging this expression and estimating the noise $\boldsymbol{\epsilon}$ using the model $\phi$

$${\hat{\boldsymbol{y}}}_{\boldsymbol{0}}=\frac{1}{\sqrt{\gamma\left( t+\Delta t,\mathbf{x} \right)}}(\boldsymbol{y}\left( t+\Delta t \right)-\sqrt{1-\gamma\left( t+\Delta t,\mathbf{x} \right)} {\hat{\boldsymbol{\epsilon}}}_{\phi}\boldsymbol{)}$$

Then, to sample from the reverse process, we can sample from the forward process, starting at ${\hat{\boldsymbol{y}}}_{\boldsymbol{0}}$. Ultimately, we find that $\boldsymbol{\mu}_{\psi}\left( \mathbf{y}\left( t+\Delta t \right) \right)$can be directly computed from a noise estimate, i.e.,

$$\boldsymbol{\mu}_{\psi}\left( \mathbf{y}\left( t+\Delta t \right) \right)=\sqrt{\gamma\left( t,\mathbf{x} \right)}{\hat{\boldsymbol{y}}}_{\boldsymbol{0}}=\frac{1}{\sqrt{1-\beta_{t}}}\left( \boldsymbol{y}\left( t+\Delta t \right)-\sqrt{1-\gamma\left( t+\Delta t,\mathbf{x} \right)} {\hat{\boldsymbol{\epsilon}}}_{\phi} \right)$$

Repeated application of this process permits sampling from the reverse process over several time steps.

**OPTIMIZATION OF CVDM**

Briefly, the CVDM model uses the following objective function for optimization

$$\mathcal{L=}\mathcal{L}_{\mathrm{prior}}+\mathcal{L}_{\infty}+\alpha\mathcal{L}_{\gamma}+\mathcal{L}_{\beta}$$

The term $\mathcal{L}_{\mathrm{prior}}=D_{KL}(q(\boldsymbol{y}_{T}|\boldsymbol{y}_{0})||p(\boldsymbol{y}_{T}))$ helps to helps the forward process produce a pure Gaussian random variable. This is computed as the KL-divergence of a standard normal prior and the terminal marginal distribution of the forward process. The term $\mathcal{L}_{\infty}$is a Monte Carlo noise estimation loss:

$$\mathcal{L}_{\infty}=\frac{1}{2}E_{\epsilon\sim\mathcal{N}\left( 0,I \right),t\sim U(\left[ 0,1 \right])}\left\| \epsilon-\hat{\epsilon} \right\|_{2}^{2}$$

The last term is a constraint on $\gamma(t,\mathbf{x})$ to ensure similarity between discrete and continuous time noise schedules. As a regularization term, $\mathcal{L}_{\gamma}=$ $\left\| \gamma^{''}(t,\boldsymbol{x}) \right\|_{2}^{2}$ . The fourth and final term $\mathcal{L}_{\beta}$ enforces the exponential decay of the signal level and constrains the diffusion process at its endpoints

$$\mathcal{L}_{\beta}=E_{t\sim U(\left[ 0,1 \right])}\left[ \left\| \frac{\partial\gamma(t,\mathbf{x})}{\partial t}+\gamma(t,\mathbf{x})\beta(t,\mathbf{x}) \right\|_{2}^{2}+\left\| \gamma\left( 0,\mathbf{x} \right)-\mathbf{1} \right\|_{2}^{2} +\left\| \gamma\left( 1,\mathbf{x} \right)-\mathbf{0} \right\|_{2}^{2} \right]$$

Optimization of (1) is carried out using the hyperparameters given in Supplementary Table 1.

**SIMULATED LEARNING OF CVDM ON LOCALIZATION MICROSCOPY DATA**

A complete listing of simulation parameters for simulations presented in Figure 2 and Figure 3 of the main text can be found in Supplementary Table 2. Training data consists of low-resolution 64x64 images, setting $\sigma_{\mathbf{x}}=1.0$ in units of low-resolution pixels, assuming a 100nm pixel size. Images are generated by sampling $\rho$=100, 200, or 500 coordinate pairs $\theta$ uniformly over the image. This is followed by integration of the PSF and multiplication of $\omega_{k}$ for individual spots by the signal photon rate, drawn uniformly between [500,1000]. Final pixel values are generated by simulation from equation (1). Camera noise is simulated from a normal distribution $p(\zeta_{k}\mathcal{)=N(}o_{k},w_{k}^{2})$ where $o_{k}=100 ADU$, $w_{k}^{2}=5.0 \mathrm{ADU}^{2}$. Furthermore, KDEs $\mathbf{y}$ have dimension 256x256. For simulated datasets, target KDEs are generated using $\sigma_{\mathbf{y}}=1.0$ in units of high-resolution pixels. This results in a high-resolution pixel size of 25nm. Models used on simulation data were trained on 500 simulated low-resolution and high-resolution pairs at each density level. Each model was subsequently validated on ten batches of simulated data at an equivalent density of its training data to obtain precision, recall, and localization errors per model.

**APPLICATION OF CVDM TO EXPERIMENTAL NANORULER IMAGES**

Low-resolution images have a pixel size of 44nm and we measure $\sigma_{\mathbf{x}}=1.23$ for the experimental PSF. The corresponding KDEs are rendered with $\sigma_{\mathbf{y}}=2.0$. In addition, each 64×64 training image contains a variable number of emitters (1–20 spots) with a signal photon rate uniform in [500, 5000], spatially uniform background rate in [100, 300]. Non-uniform background signal is added to each frame with a Gaussian random field (GRF). Spacing for simulated nanorulers is also randomized by drawing spacing from a normal distribution, to avoid learning a fixed structural pattern, while maintaining a realistic spacing between emitters. Moreover, experimental images often contain dense clusters of emitters which can perturb the diffusion process during inference. We include such clusters in our simulations with a parent-child cluster process: each parent ruler spawns 1–4 children with probability of 0.3. To ensure broad generalization, we trained for 100 epochs with 200 batches per epoch, drawing a new synthetic batch each step with no repeats.

**APPLICATION OF CVDM TO EXPERIMENTAL MICROTUBULE IMAGES**

Training dataset generation for experimental data of microtubules used $\sigma_{\mathbf{x}}=1.4$ in units of low-resolution pixels and we used a signal rate drawn uniformly between [2500,5000]. Camera noise is simulated from a normal distribution using $o_{k}=100 ADU$, $w_{k}^{2}=225.0 \mathrm{ADU}^{2}$ to account for noise amplification during summation of consecutive frames.

**SUPPLEMENTARY FIGURES AND TABLES**

Supplementary Table 1. CVDM training hyperparameters for simulated learning

| Parameter | Density Sim | Nanoruler Sim | Microtubule Sim |
| --- | --- | --- | --- |
| $\alpha$ | 0.001 | 0.001 | 0.001 |
| Epochs | 50 | 100 | 50 |
| Generation time steps | 200 | 200 | 200 |
| Learning rate | ${10}^{-4}$ | ${10}^{-4}$ | ${10}^{-4}$ |
| Batch size | 2 | 2 | 2 |

Supplementary table 2. Simulation parameters

| Parameter | Density Sim | Nanoruler Sim | Microtubule Sim |
| --- | --- | --- | --- |
| # samples | 500/density | $\infty$ | 1000 |
| Spatial distribution | Uniform | Nanoruler | Uniform |
| $\boldsymbol{x}$ dims | 64x64 | 64x64 | 64x64 |
| $\boldsymbol{y}$ dims | 256x256 | 256x256 | 256x256 |
| $\sigma_{\boldsymbol{x}}$ | 1.0 | 1.23 | 1.4 |
| $\sigma_{\boldsymbol{y}}$ | 1.0 | 2.0 | 1.0 |
| $o_{k}$ [ADU] | 100 | 100 | 100 |
| $w_{k}^{2}$ [ADU^2^] | 5.0 | 24.1 | 225 |
| Signal rate [photons] | Uniform([500,1000]) | Uniform([500,5000]) | Uniform([2500,5000]) |
| BG rate [photons] | 0 | U([100, 300]) | 0 |
| Spatial BG | No | GRF | No |
| Clusters | None | Parent-Child | None |

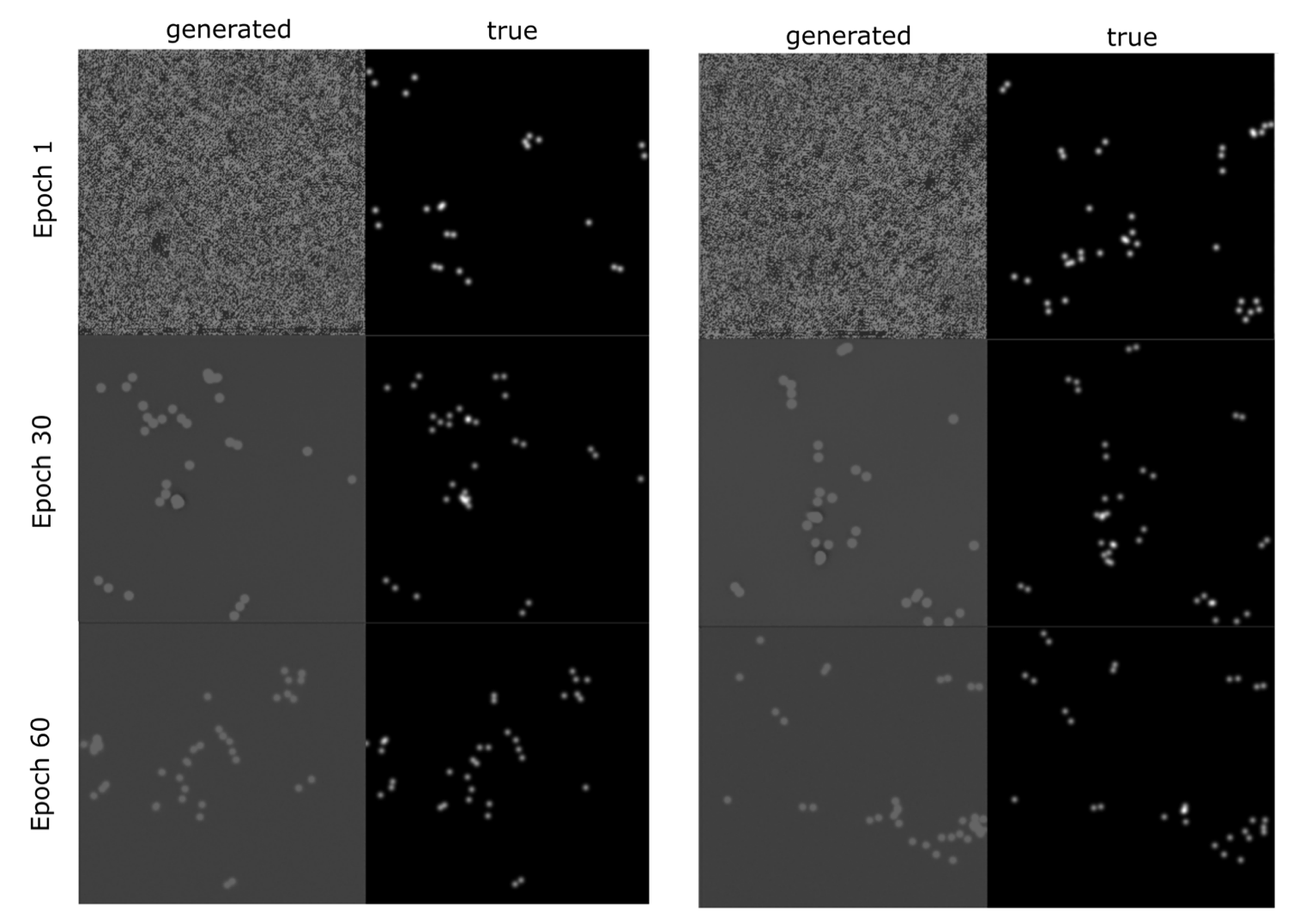

Supplementary Figure 1. Two predictions from CVDM and ground truth at epoch 1/30/60
